# Bathyarchaeota Occurrence in Shallow Marine Methane-rich Sediments (Saco do Mamanguá, Rio de Janeiro, Brazil)

**DOI:** 10.1101/2020.10.07.329656

**Authors:** R. G. Romano, A. G. Bendia, D. C. Franco, C. N. Signori, T. Yu, F. Wang, L. Jovane, V. H. Pellizari

## Abstract

Methane gas (CH_4_) reservoirs have previously been reported in tropical marine sediments of Saco do Mamanguá ria (Rio de Janeiro, Brazil). It is known that a methane microbiome can be established in methane seeps sites; however, they are still poorly characterised. In this study, we aimed to investigate the spatial and vertical distribution of the bacterial and archaeal composition and the community structure in the sediments of Saco do Mamanguá ria. For this purpose, we collected sediment samples through 100 cm long gravity corer at three different sites and performed community analysis based on 16S rRNA gene sequencing, quantification of the methyl coenzyme M reductase-encoding gene (mcrA) and geochemical analysis, including CH_4_ stable isotope. Our results revealed a biogenic trend for CH_4_ isotopic signature and a high proportion of archaeal sequences assigned as Bathyarchaeota, with a spatial distribution throughout the inner areas of the channel and the deepest strata. OTUs classified within Bathyarchaeota and Chloroflexi (Dehalococcoidia) showed positive correlation with methane concentrations, sediment depth and oxidation-reduction potential. Due to their prevalence in the microbial community, we also performed phylogenetic analysis to understand the closeness of our Bathyarchaeota OTUs with Bathyarchaeota subgroups, and the results demonstrated a close relationship particularly with Bathy-8 and Bathy-13, which possess genes for acetogenesis and methanogenesis. Although direct relations between microbial populations and the biogenic methane emissions in Saco do Mamanguá cannot be assured, our results emphasize the importance of further investigations about the potential role of Bathyarchaeota in the carbon cycling in methane-rich tropical shallow ecosystems.

## 1 INTRODUCTION

The occurrence of shallow methane (CH_4_) gas within marine sediment and the exudation points for the water column in Saco do Mamanguá ria (Rio de Janeiro, Brazil) was previously reported by Benites et al. 2015. Methane in marine sediments could have a thermogenic or a biogenic production by methanogenic Archaea through the degradation of organic debris deposited on the seafloor (Egger et al. 2018). To differ between them, it has been proposed that thermogenic methane is characterized by heavy carbon (−50 to −20‰) isotopic signatures, while biogenic methane produced locally by microbial activities is generally characterized by lighter carbon (−110 to −50‰) isotopic signatures (Whiticar 1999).

Methanogenic archaea are well known as key contributors to global CH_4_ emissions when producing gas using different sets of substrates, such as molecular hydrogen and carbon dioxide, methylated compounds and acetate (Reeburgh 2007; Stocker et al. 2013). Distinct carbon sources can be used by these methanogens (Boopathy et al. 1998), but the final enzymatic reaction that leads to the methane production is always carried out by the methyl co-enzyme M reductase (*mcrA* gene), representing a marker gene for methane-cycling archaea (Luton et al. 2002).

The main taxa involved with methanogenesis are assigned within phylum Euryarchaeota (Thauer et al. 2008), represented by the classes Methanosarcinales, Methanocaellales Methanobacteriales, Methanococcales, Methanomicrobiales, Methanopyrales (Luton et al. 2002) and Methanomassiliicoccales (Borrel et al. 2014). Novel metagenomic evidence has indicated the presence of *mcr*-enconding genes in the recently described archaeal phyla Verstraetearchaeota and Bathyarchaeota (Kelly et al. 2011; Evans et al. 2015; Vanwonterghem et al. 2016; Wang et al. 2019), however it is not clear whether they are actually involved with methane production. Metagenome-assembled genomes of Bathyarchaeota-related groups indicate a wide range of metabolic capabilities among these organisms, including the potential metabolism of methane (due to the identification of *mcr*-encoding genes), acetogenesis, and dissimilatory nitrogen and sulphur reduction. Possible interactions with anaerobic methane-oxidizing archaea, acetoclastic methanogens and heterotrophic bacteria have also been proposed for this phylum (Evans et al. 2015; Zhou et al. 2018).

To investigate the potential influence of CH_4_ concentrations on archaeal and bacterial composition and diversity, and microbial community structure at Saco do Mamanguá (Rio de Janeiro, Brazil), we combined *mcrA* gene quantification and 16S rRNA gene sequencing with geochemical and δ^13^C_CH4_ stable isotope analysis. In addition, a suite of physicochemical parameters was selected to test their influence on the microbial distribution. This study represents the first report on the spatial and vertical distribution of bacteria and archaea in a tropical methane-rich shallow water ecosystem, with emphasis on the most abundant archaeal phylum Bathyarchaeota.

## 2 MATERIALS AND METHODS

### Study sites and sampling

Geomorphologically classified as a ria (Evans and Prego 2003) Saco do Mamanguá (Rio de Janeiro, Brazil), is a gulf 11 km long and 2 km wide, oriented southwest-northeast and connected to the Atlantic Ocean, which forms a distinct environment with low river discharge (fig. 1). The average water depth of Saco do Mamanguá is approximately 5 m, and can reach a maximum of 20 m in the outer area (Rodelli et al. 2019; Brandini et al. 2019).

**Fig. 1.**
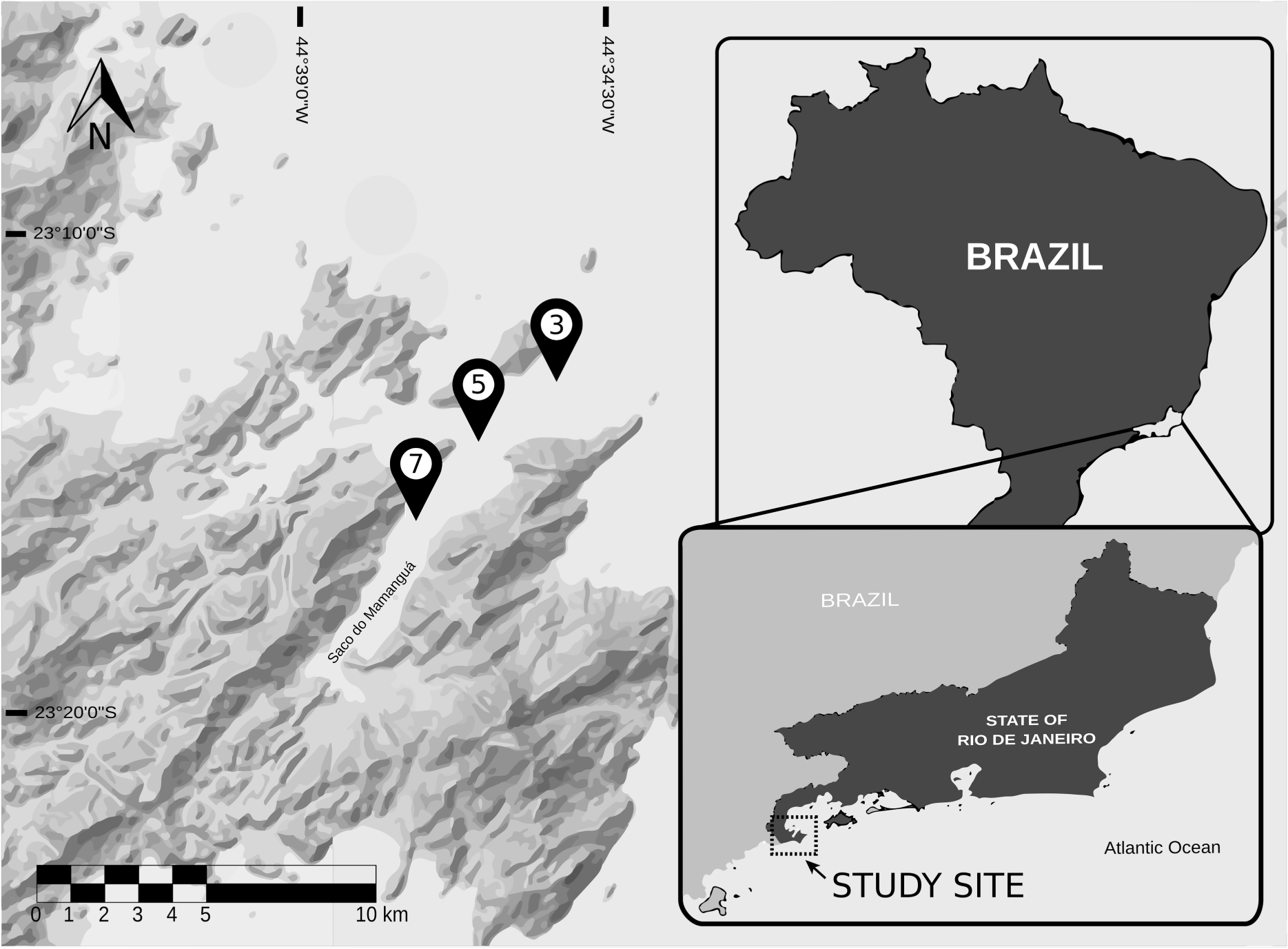
Location of Saco do Mamanguá, state of Rio de Janeiro, southeastern Brazil. The markers show the sampling sites 3, 5 and 7. The map was generated using ESRI ArcGIS.

In an oceanographic cruise onboard research vessel Alpha Delphini (owned by the Oceanographic Institute, University of São Paulo, Brazil.) in February 2015, three different sampling areas were selected, including site 3 (external area, 17.3 m water depth), site 5 (middle area, 13.2 m water depth) and site 7 (internal area, 10.4 m water depth) (fig. 1). For each sampling point, four 100cm sediment cores were sampled using a gravity corer sampler.

### Physical-chemical profile

#### Pore water chemistry and CH_4_ stable isotope

Pore water was extracted every 20 cm along one core for each sampling point, using a rhizosphere membrane (Rhizosphere Research Products B.V., The Netherlands) connected to a syringe. Dissolved oxygen, pH, electrical conductivity and redox potential were measured using a transparent glove box with a nitrogen inert atmosphere. For dissolved oxygen, a DO probe connected to a *sympHony™* (VWR®, USA) benchtop meter was used, while for pH, electric conductivity, and oxidation-reduction potential, a *sensION™* (HACH®, USA) portable meter. Measurements for CH_4_ stable isotope (δ^13^C_CH4_) were performed in the Stable Isotope Laboratory at the Geoscience Institute of the University of São Paulo, using a gas chromatography isotope ratio mass spectrometry system (Isoprime Ltd.) and following the method described by Assayag et al. (2006).

#### Methane concentration and sediment chemistry

Methane concentration was measured using a CP-4900 (Varian Inc., USA) micro-gas chromatograph equipped with a thermal conductivity detector. One core from each sampling site was cut, from top to bottom into pieces of 20 cm. At the same time, the equipment sensor was attached to the top of each strata proceeding to the measurement process.

After data acquisition, each layer of the sediment was sampled in plastic bags and sent to Laboratory of Soil Sciences, Luiz de Queiroz College of Agriculture - University of São Paulo (LSO-ESALQ) to measure the concentrations of total nitrogen, ammonium, nitrate, total sulphur, total organic matter and total organic carbon, according to previously described methods (Blume 1985; Raij et al. 2001).

### Microbial Community Profile

To determine the microbial community profiles, the sediment cores were sliced in 0-20 cm sections and stored in −20°C, using stereo plastic bags. The strata 0-20, 20-40 and 80-100 cm were selected for this study, with the exception of sampling site 3, which we did not achieve the 80-100 cm strata during sampling.

#### DNA extraction

DNA was extracted from 0.5 g of sediment using a PowerSoil DNA kit (MoBio, Carlsbad, CA, USA), according to the manufacturer’s instructions. Extracted DNA was purified using a OneStep™ PCR Inhibitor Removal kit (Zymo Research, Irvine, CA, USA) and subsequently quantified using an Qubit dsDNA HS assay (Thermo-Fisher Scientific, USA).

#### qPCR

Real-time PCR was performed using the primer pair mlas-F (GGYGGTGTMGGDTTCACMCARTA) and mcrA-R (CGTTCATBGCGTAGTTVGGRTAGT) to target the gene encoding methyl coenzyme M reductase (*mcrA*) (Kolb et al. 2003; Steinberg and Regan 2008). PCR was performed using a mixture comprising 5 μl of DNA template, 0.1 μl of bovine serum albumin (BSA; 10 mg/ml), 0.25 μl of acrylamide (10 mg/ml), 1.5 μl of each primer (1.2 µM), and 12.5 μl of SYBR GreenER qPCR SuperMix Universal (Thermo Fisher Scientific, Waltham, MA, USA), with a total volume of 25 μl. PCR was performed using a StepOne Real-Time PCR System (Thermo Fisher Scientific, USA) with an initial denaturation step at 95 °C for 10 minutes followed by 50 cycles of denaturation at 95 °C for 30 seconds, annealing at 56 °C for 45 seconds, extension at 68 °C for 45 seconds and an incubation at 81 °C for 8 seconds to allow for dissociation of potential primer dimers and nonspecific amplification products and for fluorescence detection (Freitag et al. 2010). qPCR efficiency (*E* = 10 ^(1/−slope)^) and linearity (*R*-squared value) were calculated from standard curves, and CT measurements were converted to number of copies per gram of sediment for each sample.

#### 16S rRNA gene sequencing

The V3-V4 region of the rRNA 16S gene was amplified using the universal primers 515F (GTGYCAGCMGCCGCGGTAA) and 806R (GGACTACNVGGGTWTCTAAT) (Caporaso et al. 2011), following the Earth Microbiome Project protocol (Thompson et al. 2017). Library construction and sequencing by the Illumina MiSeq 2500 (Illumina, Inc., San Diego, CA, USA) platform were performed by Macrogen Korea Laboratories (Seoul, KO).

#### Sequencing data processing and statistical analyses

The raw reads were filtered for length (> 250 bp), quality score (mean > 30), and minimum expected errors of 1.0 using the USEARCH algorithm (Edgar 2010). After the removal of chimeras using the UCHIME algorithm (Edgar et al. 2011), the remaining sequences were clustered into operational taxonomic units (OTUs) at a 97% similarity threshold. Taxonomic classification and diversity analysis were performed using the Quantitative Insights Into Microbial Ecology (QIIME) 1.8.0 pipeline (Caporaso et al. 2010). Subsequently, the singletons were removed, and taxonomic classification was performed using the SILVA 128 database (Quast et al. 2013; Yilmaz et al. 2014). The OTU table was normalized using cumulative sum scaling (CSS) for beta-diversity analysis (Paulson et al. 2013).

OTUs related to Bathyarchaeota were selected for subsequent diversity analysis. Beta-diversity was examined through UNIFRAC weighted distance analysis, which was visualized by non-metric multidimensional scaling (nMDS), with the fitting of the environmental parameters performed by applying the envfit function from the vegan package (Oksanen et al. 2010). Spearman correlations were performed to determine the relationships between environmental parameters and two different OTU groups: (1) the 30 most abundant OTUs and (2) the Bathyarchaeota-related OTUs. Statistical analyses were performed using R software (version 3.3.1), with the packages qiimer, vegan, ggplot2 flyr, and reshape2. The raw sequence data was deposited in GenBank as BioProject ID PRJNA522377.

#### Comparison of Bathyarchaeota-related OTUs with a metagenomic database

Bathyarchaeota OTUs were selected for comparison with a Bathyarchaeota database that was previously constructed using metagenome-assembled genomes by the State Key Laboratory of Microbial Metabolism (School of Life Sciences and Biotechnology, Shanghai Jiao Tong University, Shanghai, China) (Yu et al. 2017). To construct the tree, the sequences related to Bathyarchaeota were aligned using CLUSTALX 1.83. Neighbour-joining phylogenetic trees were constructed from pairwise comparisons with the Kimura 2-parameter distance model using the molecular evolutionary genetics analysis (MEGA) program, version 6 (Tamura et al. 2013). The robustness of the inferred topology was tested by 1000 bootstrap resampling.

## 3 RESULTS

### Physicochemical parameters

Physicochemical analyses showed that oxidation-reduction potential increased with depth at all sites, varying from −167.30 and 10.20 mV for the 0-20 cm layers, −239.00 and 46.00 mV for the 20-40 cm layers and −306.00 to −228.00 mV for the 80-100 cm layers. The dissolved oxygen values decreased with depth at all sampling sites, ranging from 4.24-6.96 mg/l for the 0-20 cm layers, 3.14-7.00 mg/l for the 20-40 cm layers and 1.66-3.00 mg/l for the 80-100 cm layers. Vertical distribution patterns were not observed for pH, electrical conductivity, total nitrogen, ammonium, nitrate, sulphur organic matter and organic carbon (Supplementary table A). The CH_4_ concentrations increased with depth, varying from 21.37-36.63 ppmv for the 0-20 cm layers and 118.53-377.47 ppmv for the 80-100 cm layers. The δ^13^C_CH4_ values fluctuated from −73.38 to −41.70 ‰ for site 3, −101.00 to −60.76 ‰ for site 5 and −68.85 to −52.67 ‰ for site 7 (Supplementary table A).

#### Quantification of the mcrA gene

For all sites, the *mcrA* gene copy numbers were higher in the 0-20 cm interval than those observed in the other strata, ranging from 1.5 × 105 to 1.0 × 106 copies/g of sediment (fig. 2).

**Fig. 2.**
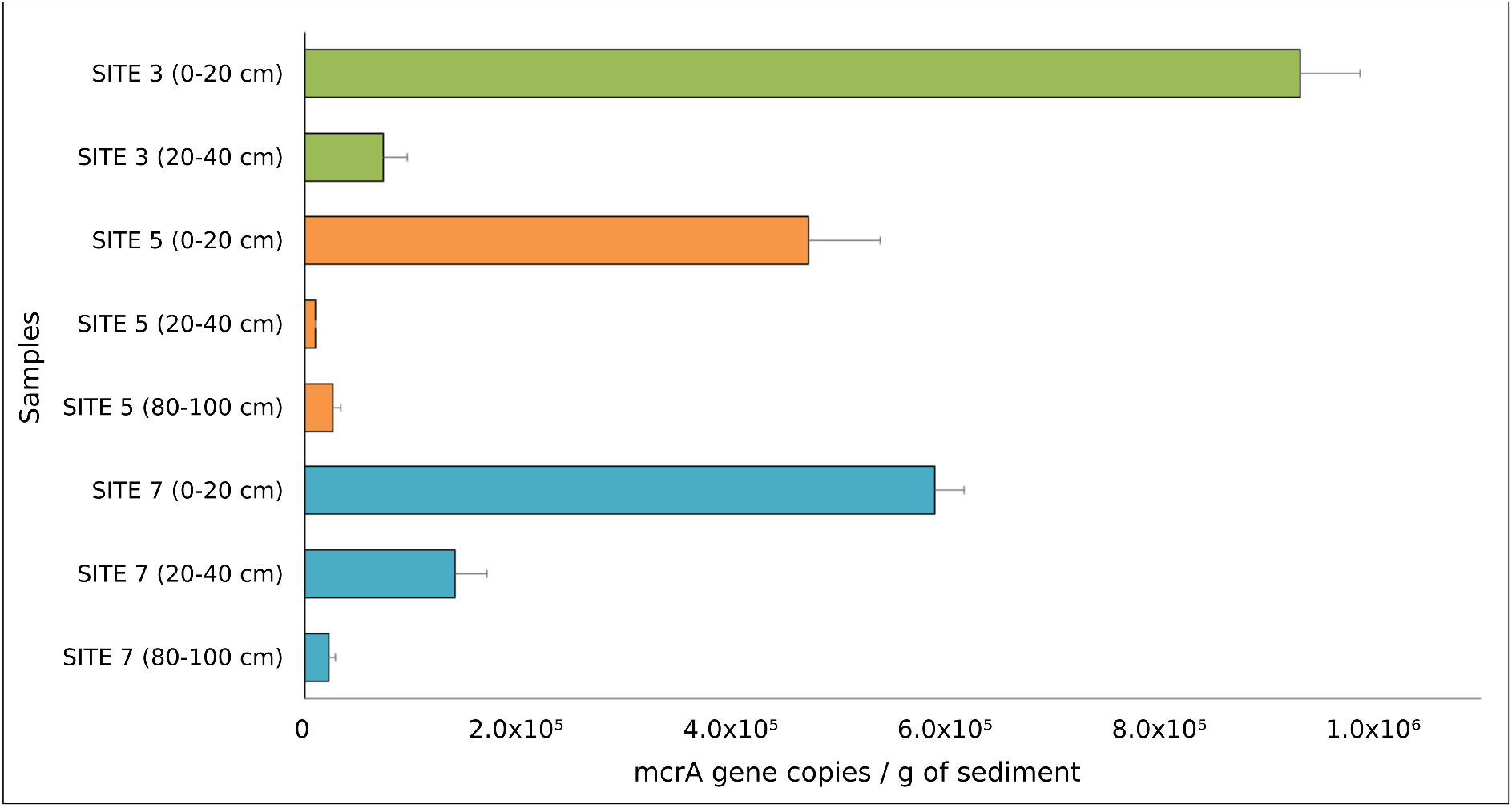
qPCR results for the mcrA gene copies/g of sediment from Saco do Mamanguá. The lines at the end of the bars show the standard deviation, and each colour represents a different sampling site (green = sampling site 3, orange = site 5 and blue = site 7).

### Microbial community structure

Through 16S rRNA gene sequencing, a total of 1,748,277 high-quality reads comprising 10,817 OTUs were obtained from the eight sediment samples from Saco do Mamanguá. For the total microbial community at the phylum level (fig. 3a), the relatively abundant phyla in the surface layers were Proteobacteria (30 %), Chloroflexi (17.7 %), Planctomycetes (8.1 %) and Acidobacteria (7.0 %). In the deepest layers, the predominant phyla were Chloroflexi (33-17.5 %), Bathyarchaeota (18.7-8.7 %), Aminicenantes (9.0-5.8 %), and unclassified phyla (1.2-3.7 %).

**Fig. 3.**
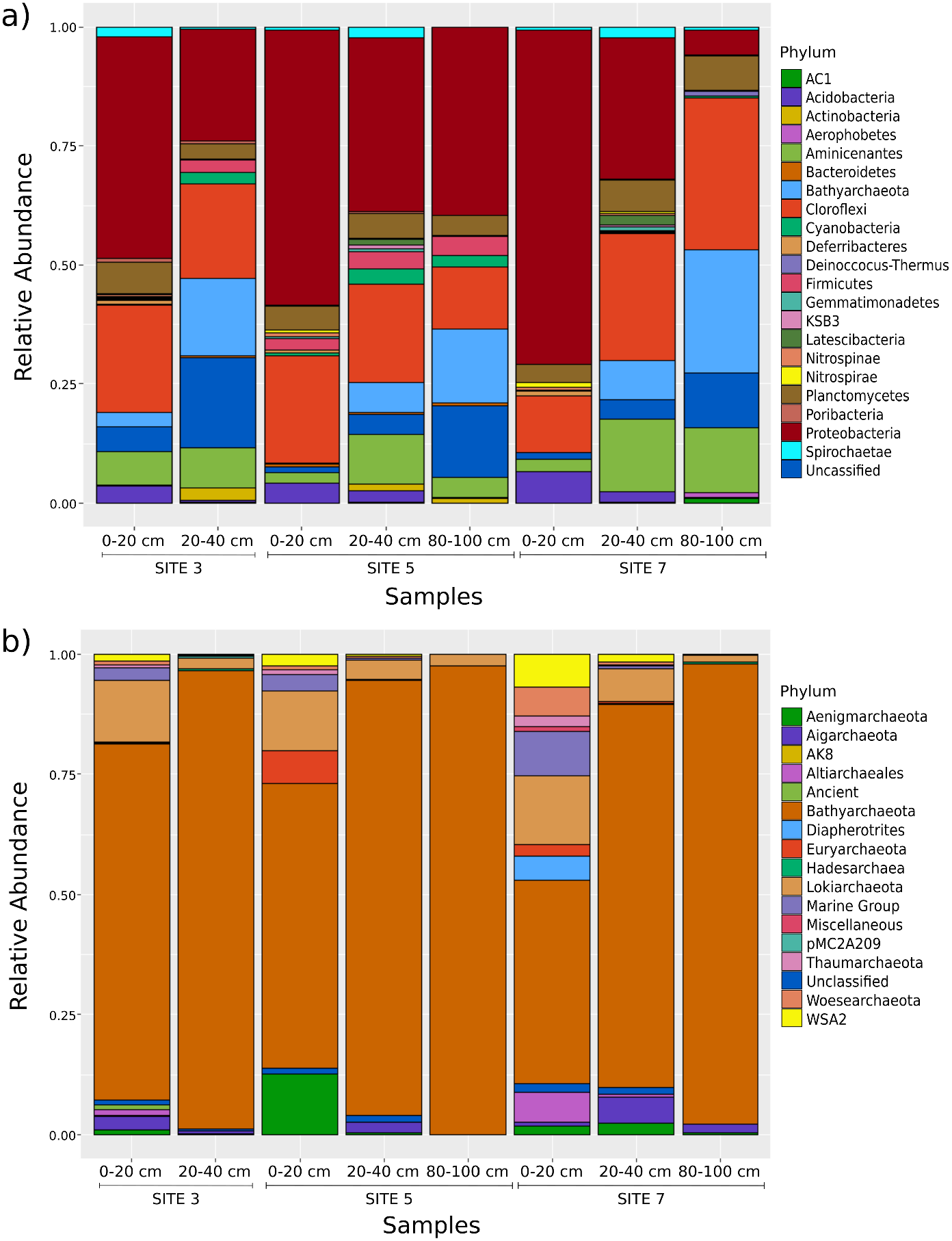
Relative abundance (%) of OTUs from (a) dominant (>0.1%) bacterial and archaeal phylotypes and (b) total archaeal phylotypes. The colours of the bars and the corresponding phyla are listed on the right. The relative abundances and samples are shown on the y- and x -axes, respectively.

With regard to the archaeal communities (fig. 3b), the phylum Bathyarchaeota accounted for 79.3% of the total detected OTUs, whose relative proportion increased with depth. In relation to the total archaeal OTUs, the relative abundance of Bathyarchaeota was 74.0% (0-20 cm) and 95.4% (20-40 cm) for site 3; 59.3% (0-20 cm), 90.5% (20-40 cm) and 97.4% (80-100 cm) for site 5; and 42.4% (0-20 cm), 79.6% (20-40 cm) and 95.6% (80-100 cm) for site 7. The phylum Euryarchaeota represented only 1.3% of the total archaeal community, and its proportion decreased with depth, accounting for 0.3% (0-20 cm and 20-40 cm) of the communities at site 3; 6.8% (0-20 cm), 0.1% (20-40 cm) and 0.0% (80-100 cm) of the communities at in site 5; and 2.4% (0-20 cm), 0.4% (20-40 cm) and 0.0% (80-100 cm) of the communities at in site 7. In general, all sediment cores showed a decrease in richness and alpha-diversity with depth (Supplementary table B).

### Environmental parameters and microbial communities

We performed nMDS statistical analyses to verify the influences of the main environmental drivers and CH_4_ gradients on the total microbial community and Bathyarchaeota diversity. The non-metric multidimensional scaling (fig. 4a), revealed that only the samples from surface layers were grouped closely and that the sediment depth, organic matter and organic carbon (*p*-values = 0.077, 0.06 and 0.065, respectively) were significantly correlated (*p* < 0.1). When considering only the Bathyarchaeota community (fig. 4b), significant correlations were observed with CH_4_, sediment depth and oxidation-reduction potential (*p*-values = 0.011, 0.062 and 0.011, respectively). The CH_4_ concentrations observed between the sediment layers, influenced the Bathyarchaeota community only at the deepest layers

**Fig. 4.**
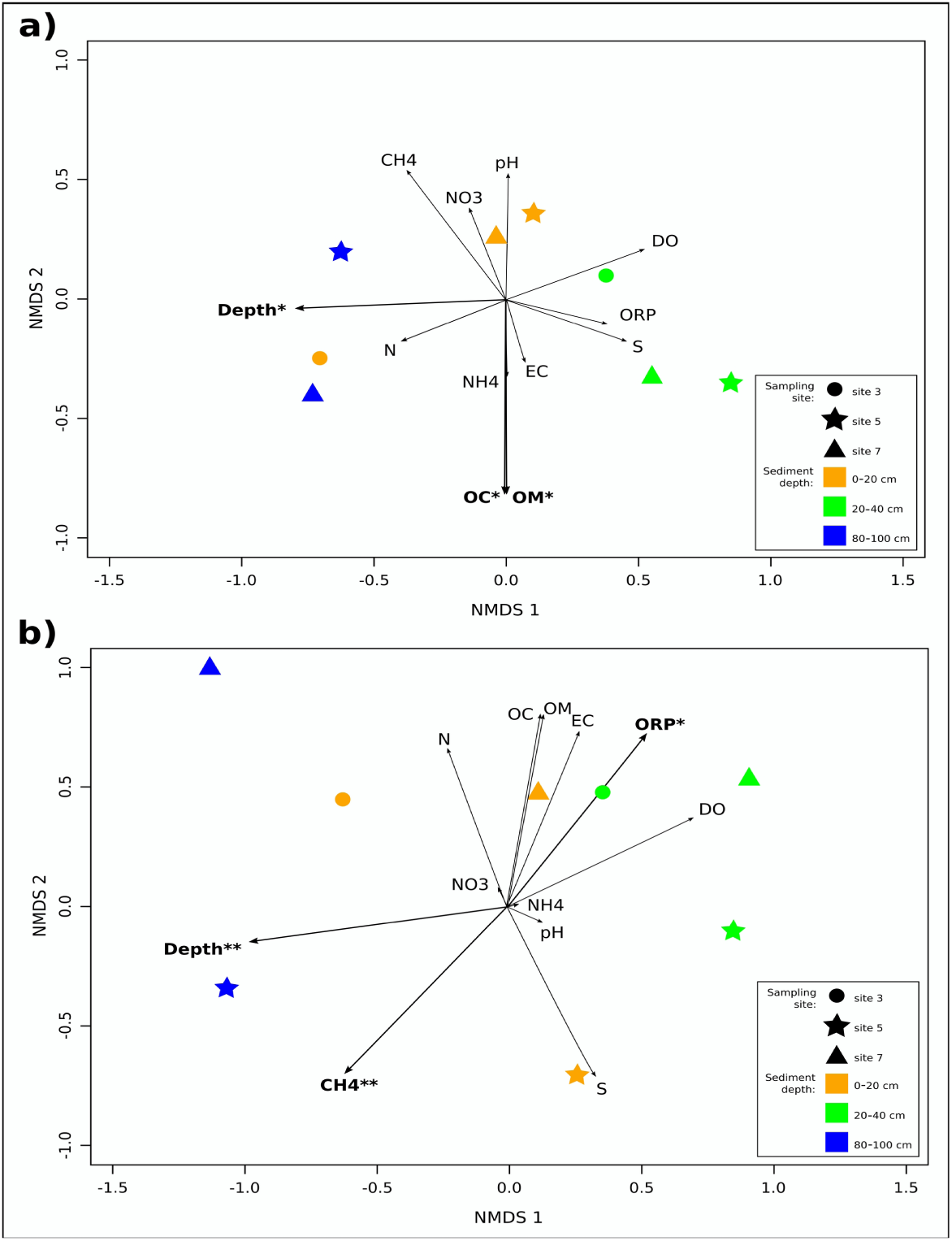
Non-metric multidimensional scaling (nMDS) ordination based on weighted UNIFRAC distance, with plotting of the environmental parameters for all OTUs (a) and for the Bathyarchaeota phylum OTUs alone (b). The arrows represents the direction and strength of the environmental parameter (Depth = depth of the sediment layer, DO = dissolved oxygen, pH = potential for hydrogen, ORP = oxidation-reduction potential, EC = electrical conductivity, N = total nitrogen, NH4 = ammonium, NO3 = nitrate, S = total sulphur, OM = total organic matter, OC = total organic carbon and CH4 = methane). Parameters in bold represent significant correlation (envfit, * = p < 0.1 and ** = p < 0.05) with stress values= 0.0647 (a) and 0.186 (b).

Additionally, we determined Spearman’s correlation coefficients to understand the relationship between environmental drivers and the 30 most abundant OTUs and only with Bathyarchaeota. Most proteobacterial OTUs presented significant negative correlations with the sediment depth, while only OTU19 (assigned as BD7-8 Marine Group) showed a positive correlation with the dissolved oxygen. Bathyarchaeota comprised four of the 30 most abundant OTUs, showing the same strong positive correlation profile with the sediment depth and CH _4_ concentrations. In addition, two Dehalococcoidia OTUs were positively associated with CH_4_ (fig. 5a). Of the 10 Bathyarchaeota OTUs detected in our study (fig. 5b), eight exhibited a positive correlation with CH _4_ and depth, including the six most abundant OTUs. Other Bathyarchaeota OTUs exhibited significantly negative correlations with organic carbon, organic matter and dissolved oxygen. The *p*-values obtained for the correlations are shown in supplementary tables C and D.

**Fig. 5.**
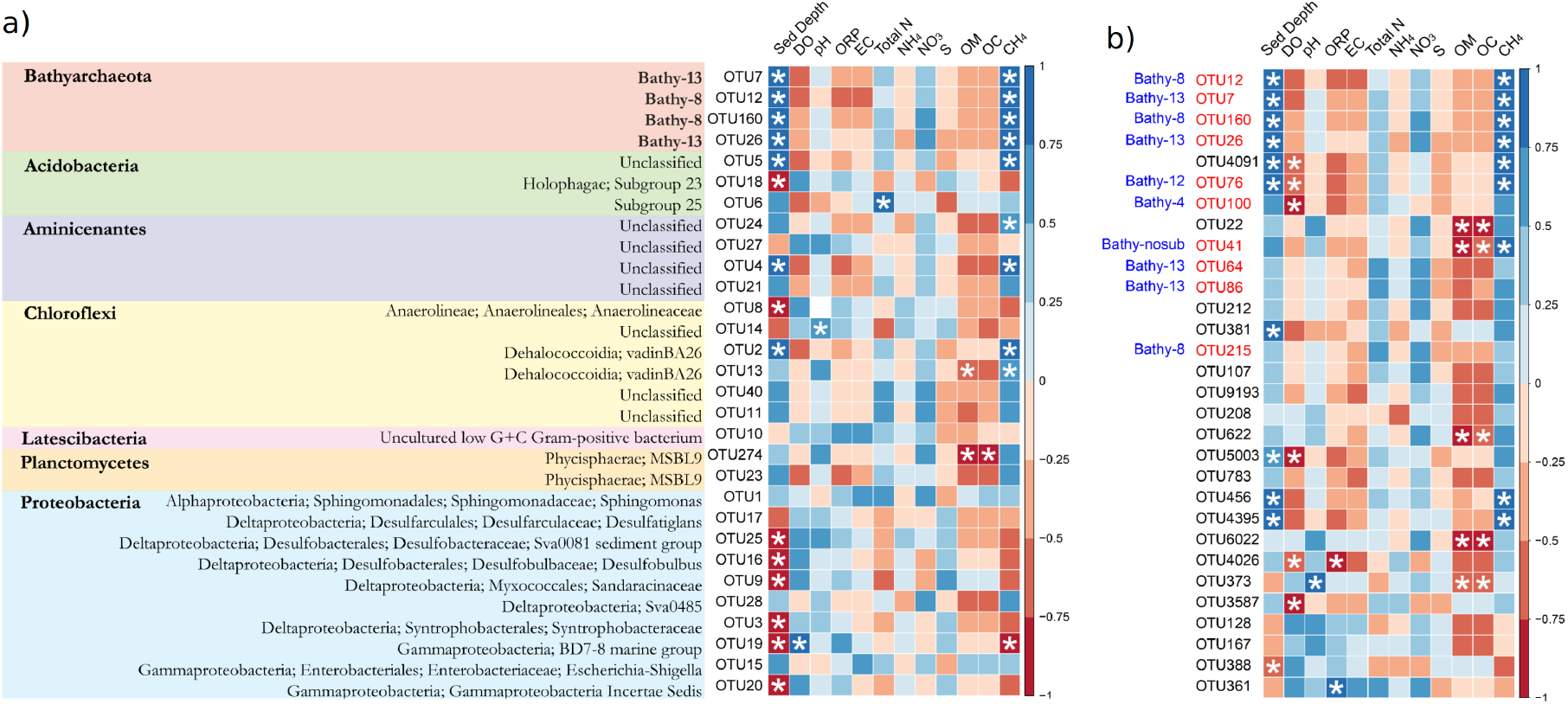
Spearman correlation (a) between the 30 most abundant OTUs and the physicochemical parameters with the highest rated taxonomic level and (b) between the Bathyarchaeota OTUs and physicochemical parameters with the corresponding Bathyarchaeota subgroups in blue. OTUs are organized by abundance in descending order from top to bottom. The environmental parameters are designated as Sed. Depth (depth of the sediment layer), DO (dissolved oxygen), pH (potential for hydrogen), ORP (oxidation-reduction potential), EC (electrical conductivity), Total N (total nitrogen), NH4 (ammonium), NO3 (nitrate), S (total sulphur), OM (total organic matter), OC (total organic carbon) and CH4 (methane). The names in red correspond to Bathyarchaeota OTU and those in blue to its associated subgroup from an external database. A white asterisk marks significant correlation.

### Phylogenetic relationships between Bathyarchaeota OTUs

The 10 Bathyarchaeota OTUs obtained in this study were assigned to five Bathyarchaeota subgroups (fig. 6). OTUs 7, 26, 64 and 86 are phylogenetically associated with Bathy-13, while OTUs 12, 160 and 125 are related to subgroup Bathy-8. OTUs 76, 100 and 41 were phylogenetically related to Bathy-12, Bathy-4 and Bathy-10, respectively.

**Fig. 6.**
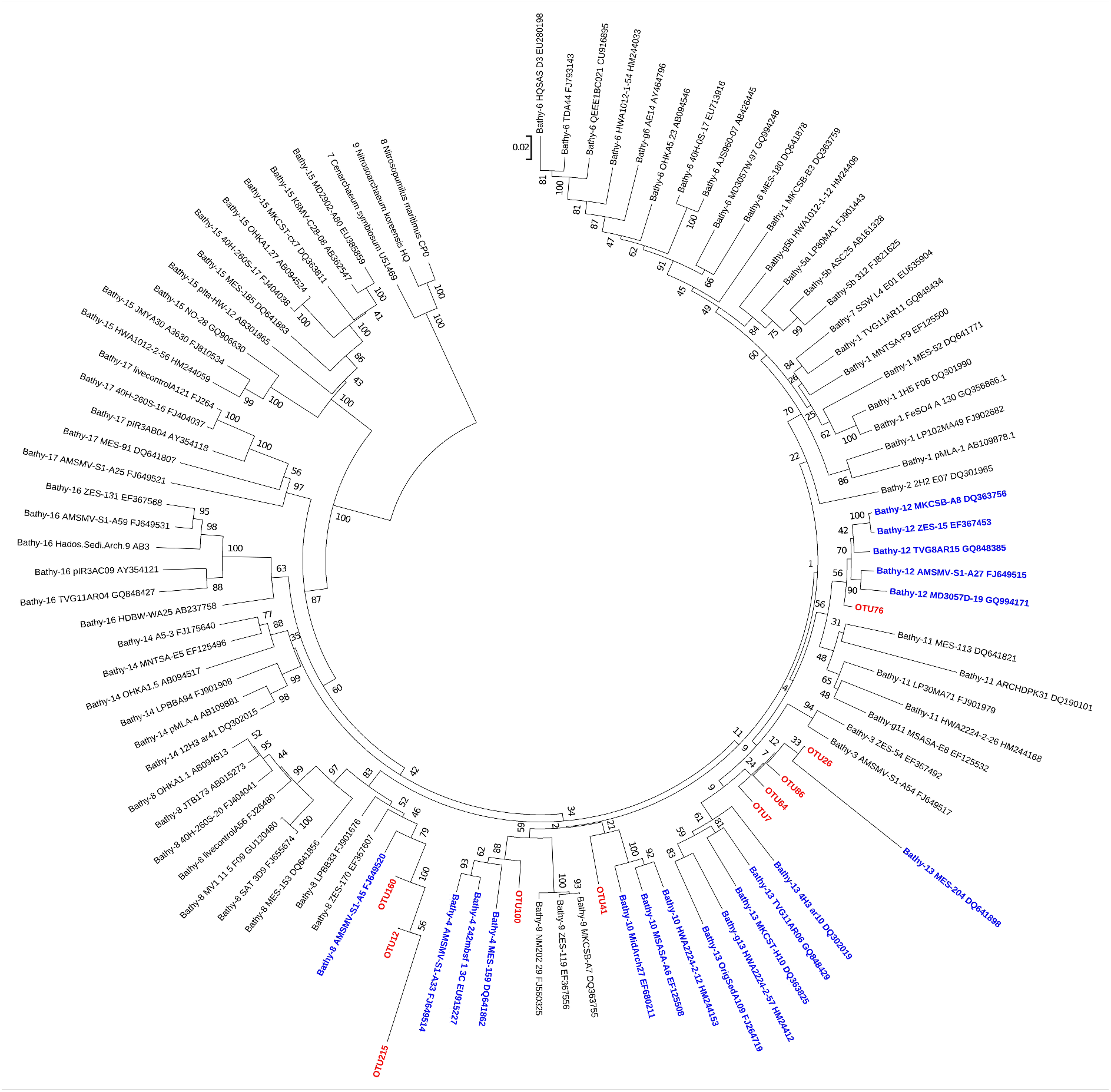
Neighbour-joining (NJ) phylogenetic tree based on 16S rRNA gene sequences of Bathyarchaeota obtained from Illumina sequencing data. The 16S rRNA genes from Cenarchaeum symbiosum (U51469), Nitrosopumilus maritimus (CP000866) and Nitrosoarchaeum koreensis (HQ331116) were used as an outgroup. The robustness of the inferred topology was tested by 1000 bootstrap resamplings. Scale bar, 0.02 substitutions per nucleotide position. In red are OTUs corresponding to Bathyarchaeota described in this work and in blue, to their associated subgroup.

## 4 DISCUSSION

Based on carbon and hydrogen isotope information from methane in natural environments, Whiticar 1999 proposed that δ^13^C_CH4_ ranging from −110 to −50 ‰ correspond to a biogenic origin of CH_4_, being the values between −110 to −60‰ representing CH_4_ highly depleted in ^13^C (methane production pathway via CO_2_ reduction) and −65 to −50‰ methane more enriched in ^13^C (acetoclastic fermentation). All of the isotope results that we obtained in this study fall within this range (except for the site 3 0-20 cm sample, which was slightly above), indicating a trend for biogenic CH_4_ origin in Saco do Mamanguá sediments. Little is known about this topic in tropical marine shallow sediments; however, biogenic signatures from CH_4_ gas exudation were also registered in fjords and freshwater environments, such as shallow marine sediments from fjords in Norway (Sauer et al. 2016), freshwater sediments from south-eastern Poland (Gruca-Rokosz and Koszelnik 2018) and tropical floodplain river in Brazil (Ballester and Santos 2001).

Our samples harboured bacterial and archaeal phyla that are typically present in shallow and deep marine sediments, such as members of Proteobacteria, Bacteroidetes, Actinobacteria, Chloroflexi, and Acidobacteria (Petro et al. 2017; Nair et al. 2017). Nevertheless, the majority of archaeal sequences were assigned to the phylum Bathyarchaeota, while Euryarchaeota constituted a negligible portion of the total abundance, which is in disagreement with the results presented in the literature for methane production in marine sediments (Gribaldo and Brochier-Armanet 2006; Lever 2016; Vanwonterghem et al. 2016; Wang et al. 2019). Although the results corroborate with the main archaeal phyla described for shallow marine sediments (e.g., Bathyarchaeota, Crenarchaeota, Euryarchaeota and Thaumarchaeota) (Ruff et al. 2016; Pan et al. 2019), a different distribution was observed from that reported for tropical environments, where the occurrence of Bathyarchaeota was much higher in comparison with Euryarchaeota.

With respect to its geographical distribution in the studied area, the occurrence of Bathyarchaeota seems to be greater at the inner area of the channel and its abundance increases with depth. This distribution follows patterns previously observed for CH_4_, in which (1) the occurrence of CH_4_ along the channel decreases towards the open ocean (Benites et al. 2015); and (2) the concentration of CH_4_ in sediments increases with depth, as observed by our results of relative gas composition. Our study showed that several Bathyarchaeota OTUs were strongly correlated with the presence of CH_4_ through Spearman correlation and ordination analysis using the *envfit* function. In addition, the Dehalococcoidia OTUs also showed a positive correlation with CH_4_, suggesting a potential competitive or syntrophic relationship with methanogenic Archaea, as observed in previous studies (e.g., Aulenta et al., 2007; Men et al., 2012; Wen et al., 2015). Petro et al. 2017, observed the co-occurrence of Dehalococcoidia and Bathyarchaeota in shallow sediments from Aarhus Bay (Denmark), although their ecological relationship remains unknown. In this study, we observed a relatively low abundance of Euryarchaeota and only in the superficial strata of the outer sampling area. This distribution of Euryarchaeota does not follow the trend of CH_4_ concentration, and there were no significant Spearman correlations between this phylum and CH_4_ concentration. These results indicate that: (1) Euryarchaeota is probably not leading the current production of biogenic CH_4_ in Saco do Mamanguá sediments or (2) methane can probably be produced deeper in sediments by euryarchaeotal communities and diffused in shallow sediments.

As microbial methanogenesis has been primarily described in anaerobic environments, it is expected that its occurrence will increase with increasing depth in marine sediments (Wilms et al. 2007). However, our qPCR results showed an atypical *mcrA* distribution, where greater gene abundance was observed at the surface and decreased with depth. Although a previous study (Evans et al. 2015) have observed the presence of non-euryarchaeotal MCR-encoding genes in the genomes of Bathyarchaeota, it is likely that the *mcrA* primers we used only targeted Euryarchaeota *mcrA* genes, which would explain the Euryarchaeota/Bathyarchaeota trend for depth observed by 16S rRNA sequencing.

By comparing our Bathyarchaeota-related OTUs with a metagenomic database, we identified four OTUs associated with the subgroup Bathy-13, which was previously described as a new phylotype that is potentially related to acetogenesis (Lloyd et al. 2013; Lazar et al. 2016; He et al. 2016; Yu et al. 2018; Feng et al. 2019). In addition, three OTUs were related to the subgroup Bathy-8 (Table 1), members of which have been described as potential acetogens and methanogens due to the detection of genes associated with these pathways (as MCR encoders) in metagenome-assembled genomes. However, it is not possible to confirm whether they are involved with acetate or methane production, and more evidence is needed to understand their potential roles in these processes (Evans et al. 2015; Feng et al. 2019). In addition, the OTUs related to Bathy-8 detected in this study are among the most abundant bathyarchaeotal OTUs, with the majority being positively correlated with the CH_4_ concentrations in Saco do Mamanguá sediments.

**Table 1.** Correspondence of the Bathyarchaeota OTUs obtained in this work with the assigned Bathyarchaeota subgroups. The third column shows the potentially related metabolic potentials suggested by the presence of these pathways in the metagenome-assembled genomes revealed by previous studies (Feng et al., 2019; He et al., 2016; Lazar et al., 2016; Lloyd et al., 2013; Yu et al., 2018).

| OTU | Corresponding Bathyarchaeota Subgroup | Potential Related Metabolism |
| --- | --- | --- |
| OTU7 | Bathy-13 | Acetogenesis |
| OTU26 |  |  |
| OTU64 |  |  |
| OTU86 |  |  |
| OTU12 | Bathy-8 | Acetogenesis/Methanogenesis |
| OTU160 |  |  |
| OTU215 |  |  |
| OTU76 | Bathy-12 | Undetermined |
| OTU100 | Bathy-4 |  |
| OTU41 | Bathy-10 |  |

New evidence has shown that Bathy-8 members are capable of organoautotrophic growth using lignin as an energy source (Yu et al. 2018), demonstrating their important role in the degradation of terrestrial-derived lignin in shallow and anoxic marine sediments. In fact, Benites et al. (2015), noted that buried vegetation in Saco do Mamanguá can provide an important source of organic matter for methanogenic processes. Thus, the recent findings of Yu et al. (2018), should drive further investigation of Bathyarchaeota in Saco do Mamanguá sediments to assess their role in anaerobic degradation of lignin and its potential relationship with CH_4_ production.

This study provides the first report on the spatial and vertical distribution of bacteria and archaea in the sediments of the Brazilian coastal ria and emphasizes the dominance of Bathyarchaeota over other archaeal phyla in an environment likely to be rich in biogenic methane. While the total community was influenced by the sediment depth, organic matter and carbon, the Bathyarchaeota community was driven by the sediment depth in addition to CH_4_ concentrations and oxidation-reduction potential. We identified the subgroups Bathy-8 and Bathy-13 as the most abundant Bathyarchaeota, distributed mainly along the inner area of the channel and in the deepest strata and harbouring genes for methanogenesis and acetogenesis, although their role in these metabolisms is still unclear. Our findings encourage further efforts to elucidate the ecological role of this phylum in the carbon cycle of methane-rich tropical coastal ecosystems.

## Acknowledgments

We are grateful to the Centro Oceanográfico de Registros Estratigráficos (CORE) laboratory (Oceanographic Institute - University of São Paulo) and M.S. Rosa Gamba and M.S. Natascha Bergo for their scientific support.

## 7 SUPPLEMENTARY MATERIAL

**Supplementary table A.** Environmental parameters of sediment samples from Saco do Mamanguá. The parameters and corresponding units are designated as SD (sediment depth, cm), DO (dissolved oxygen, mg.l^-1^), pH (potential of hydrogen), ORP (oxidation-reduction potential, mV), EC (electric conductivity, ms.cm^-1^), N (total nitrogen, mg.kg^-1^), NH_4_^+^ (ammonium, mg.kg^-1^), NO_3_^-^ (nitrate, mg.kg^-1^), S (total sulphur, mg.dm^-3^), OM (total organic matter, g.kg^-1^), OC (total organic carbon, g.kg^-1^), CH_4_ (methane, mg.dm^-3^) and δ^13^C_CH4_ (stable isotope, ‰).

| Sample | SD | DO | pH | ORP | EC | N | NH<sub>4</sub><sup>+</sup> | NO<sub>3</sub><sup>-</sup> | S | OM | OC | CH<sub>4</sub> | δ<sup>13</sup>C<sub>CH4</sub> |
| --- | --- | --- | --- | --- | --- | --- | --- | --- | --- | --- | --- | --- | --- |
| Site 3 0-20<br>cm | 20 | 4.24 | 7.77 | -55.6 | 52 | 1120 | 123 | 46 | 776 | 47 | 27 | 25.89 | -73.38 |
| Site 3 20-40<br>cm | 40 | 3.56 | 7.68 | -97.9 | 51.8 | 1190 | 123 | 102 | 1474 | 73 | 42 | 47.09 | -41.7 |
| Site 5 0-20<br>cm | 20 | 6.96 | 7.78 | 10.2 | 56.1 | 560 | 130 | 67 | 2176 | 66 | 38 | 36.63 | -91.72 |
| Site 5 20-40<br>cm | 40 | 7 | 7.72 | 46 | 55.4 | 763 | 53 | 123 | 1392 | 65 | 38 | 71.79 | -101 |
| Site 5 80-100<br>cm | 100 | 1.66 | 7.67 | -228 | 54.2 | 707 | 105 | 35 | 1162 | 77 | 45 | 118.53 | -60.76 |
| Site 7 0-20<br>cm | 20 | 4.4 | 7.55 | -167.3 | 51.6 | 280 | 95 | 39 | 2753 | 76 | 44 | 21.37 | -52.67 |
| Site 7 20-40<br>cm | 40 | 3.14 | 7.8 | -239 | 51.7 | 252 | 112 | 60 | 2285 | 28 | 16 | 137.5 | -49.43 |
| Site 7 80-100<br>cm | 100 | 3 | 7.69 | -306 | 50.9 | 630 | 105 | 63 | 1825 | 35 | 20 | 377.47 | -68.85 |

**Supplementary table B.**
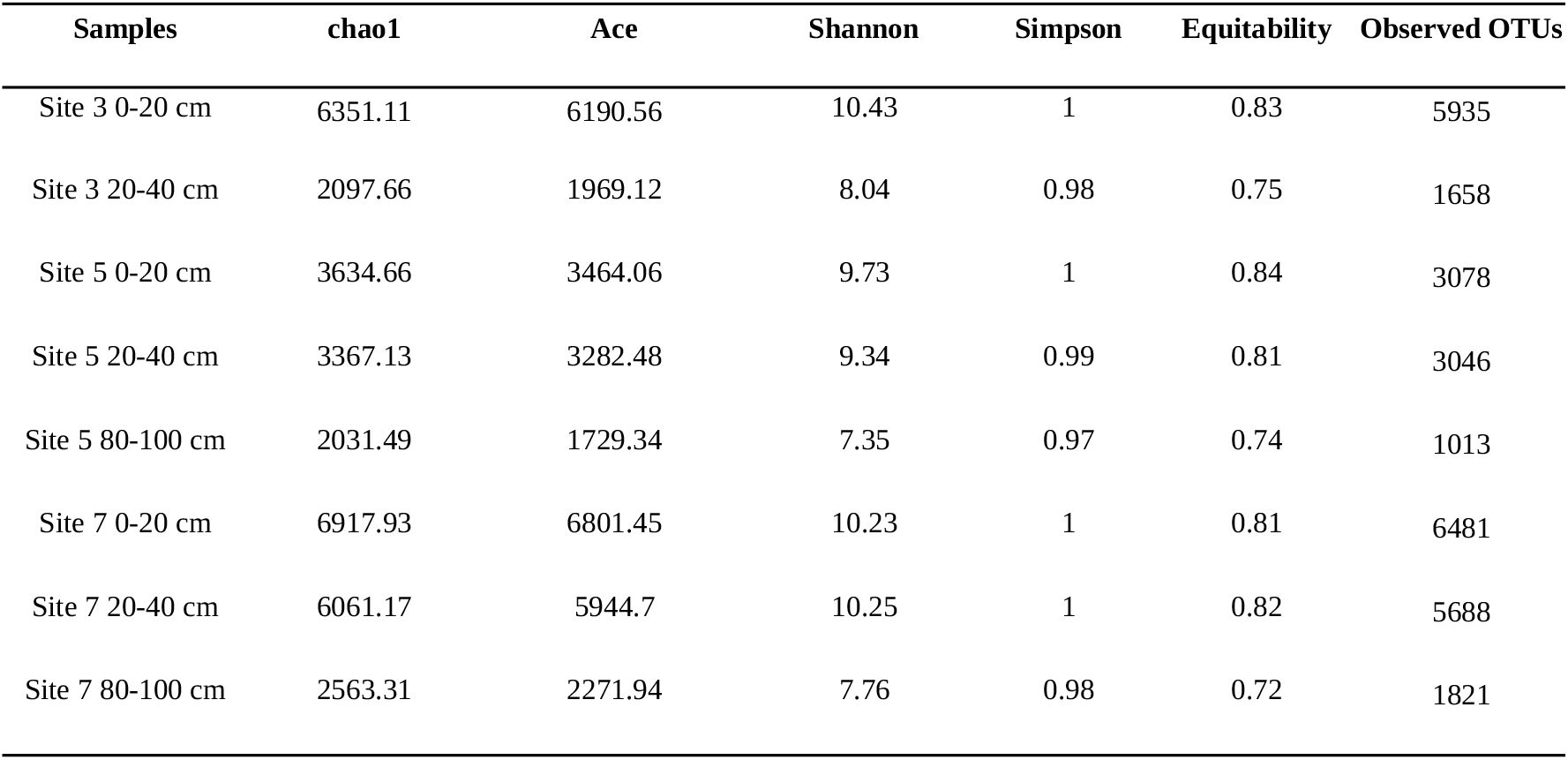
Richness and alpha diversity indices of Bacteria and Archaea from Saco do Mamanguá, using the QIIME 1.8.0 pipeline.

**Supplementary table C.**
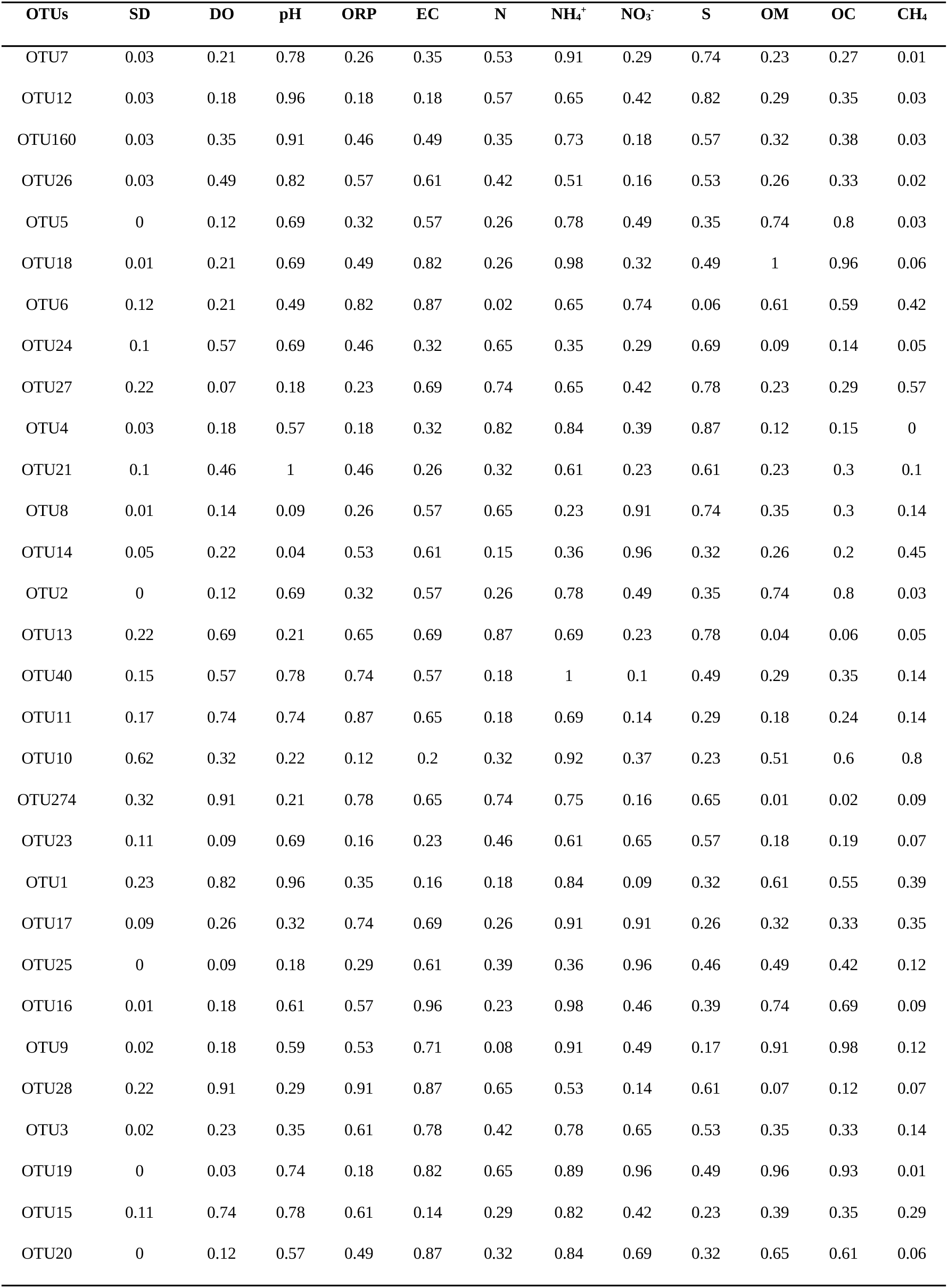
P-values for Spearman correlations between the 30 most relatively abundant OTUs and environmental parameters. Correlations were considered statistically significant when p-value<0.05. The parameters are designated as SD (sediment depth), DO (dissolved oxygen), pH (potential of hydrogen), ORP (oxidation-reduction potential), EC (electric conductivity), N (total nitrogen), H_4_^+^ (ammonium), NO_3_^-^ (nitrate), S (total sulphur), OM (total organic matter), OC (total organic carbon) and CH_4_ (methane).

| OTUs | SD | DO | pH | ORP | EC | N | $\text{NH}_4^+$ | $\text{NO}_3^-$ | S | OM | OC | $\text{CH}_4$ |
| --- | --- | --- | --- | --- | --- | --- | --- | --- | --- | --- | --- | --- |
| OTU7 | 0.03 | 0.21 | 0.78 | 0.26 | 0.35 | 0.53 | 0.91 | 0.29 | 0.74 | 0.23 | 0.27 | 0.01 |
| OTU12 | 0.03 | 0.18 | 0.96 | 0.18 | 0.18 | 0.57 | 0.65 | 0.42 | 0.82 | 0.29 | 0.35 | 0.03 |
| OTU160 | 0.03 | 0.35 | 0.91 | 0.46 | 0.49 | 0.35 | 0.73 | 0.18 | 0.57 | 0.32 | 0.38 | 0.03 |
| OTU26 | 0.03 | 0.49 | 0.82 | 0.57 | 0.61 | 0.42 | 0.51 | 0.16 | 0.53 | 0.26 | 0.33 | 0.02 |
| OTU5 | 0 | 0.12 | 0.69 | 0.32 | 0.57 | 0.26 | 0.78 | 0.49 | 0.35 | 0.74 | 0.8 | 0.03 |
| OTU18 | 0.01 | 0.21 | 0.69 | 0.49 | 0.82 | 0.26 | 0.98 | 0.32 | 0.49 | 1 | 0.96 | 0.06 |
| OTU6 | 0.12 | 0.21 | 0.49 | 0.82 | 0.87 | 0.02 | 0.65 | 0.74 | 0.06 | 0.61 | 0.59 | 0.42 |
| OTU24 | 0.1 | 0.57 | 0.69 | 0.46 | 0.32 | 0.65 | 0.35 | 0.29 | 0.69 | 0.09 | 0.14 | 0.05 |
| OTU27 | 0.22 | 0.07 | 0.18 | 0.23 | 0.69 | 0.74 | 0.65 | 0.42 | 0.78 | 0.23 | 0.29 | 0.57 |
| OTU4 | 0.03 | 0.18 | 0.57 | 0.18 | 0.32 | 0.82 | 0.84 | 0.39 | 0.87 | 0.12 | 0.15 | 0 |
| OTU21 | 0.1 | 0.46 | 1 | 0.46 | 0.26 | 0.32 | 0.61 | 0.23 | 0.61 | 0.23 | 0.3 | 0.1 |
| OTU8 | 0.01 | 0.14 | 0.09 | 0.26 | 0.57 | 0.65 | 0.23 | 0.91 | 0.74 | 0.35 | 0.3 | 0.14 |
| OTU14 | 0.05 | 0.22 | 0.04 | 0.53 | 0.61 | 0.15 | 0.36 | 0.96 | 0.32 | 0.26 | 0.2 | 0.45 |
| OTU2 | 0 | 0.12 | 0.69 | 0.32 | 0.57 | 0.26 | 0.78 | 0.49 | 0.35 | 0.74 | 0.8 | 0.03 |
| OTU13 | 0.22 | 0.69 | 0.21 | 0.65 | 0.69 | 0.87 | 0.69 | 0.23 | 0.78 | 0.04 | 0.06 | 0.05 |
| OTU40 | 0.15 | 0.57 | 0.78 | 0.74 | 0.57 | 0.18 | 1 | 0.1 | 0.49 | 0.29 | 0.35 | 0.14 |
| OTU11 | 0.17 | 0.74 | 0.74 | 0.87 | 0.65 | 0.18 | 0.69 | 0.14 | 0.29 | 0.18 | 0.24 | 0.14 |
| OTU10 | 0.62 | 0.32 | 0.22 | 0.12 | 0.2 | 0.32 | 0.92 | 0.37 | 0.23 | 0.51 | 0.6 | 0.8 |
| OTU274 | 0.32 | 0.91 | 0.21 | 0.78 | 0.65 | 0.74 | 0.75 | 0.16 | 0.65 | 0.01 | 0.02 | 0.09 |
| OTU23 | 0.11 | 0.09 | 0.69 | 0.16 | 0.23 | 0.46 | 0.61 | 0.65 | 0.57 | 0.18 | 0.19 | 0.07 |
| OTU1 | 0.23 | 0.82 | 0.96 | 0.35 | 0.16 | 0.18 | 0.84 | 0.09 | 0.32 | 0.61 | 0.55 | 0.39 |
| OTU17 | 0.09 | 0.26 | 0.32 | 0.74 | 0.69 | 0.26 | 0.91 | 0.91 | 0.26 | 0.32 | 0.33 | 0.35 |
| OTU25 | 0 | 0.09 | 0.18 | 0.29 | 0.61 | 0.39 | 0.36 | 0.96 | 0.46 | 0.49 | 0.42 | 0.12 |
| OTU16 | 0.01 | 0.18 | 0.61 | 0.57 | 0.96 | 0.23 | 0.98 | 0.46 | 0.39 | 0.74 | 0.69 | 0.09 |
| OTU9 | 0.02 | 0.18 | 0.59 | 0.53 | 0.71 | 0.08 | 0.91 | 0.49 | 0.17 | 0.91 | 0.98 | 0.12 |
| OTU28 | 0.22 | 0.91 | 0.29 | 0.91 | 0.87 | 0.65 | 0.53 | 0.14 | 0.61 | 0.07 | 0.12 | 0.07 |
| OTU3 | 0.02 | 0.23 | 0.35 | 0.61 | 0.78 | 0.42 | 0.78 | 0.65 | 0.53 | 0.35 | 0.33 | 0.14 |
| OTU19 | 0 | 0.03 | 0.74 | 0.18 | 0.82 | 0.65 | 0.89 | 0.96 | 0.49 | 0.96 | 0.93 | 0.01 |
| OTU15 | 0.11 | 0.74 | 0.78 | 0.61 | 0.14 | 0.29 | 0.82 | 0.42 | 0.23 | 0.39 | 0.35 | 0.29 |
| OTU20 | 0 | 0.12 | 0.57 | 0.49 | 0.87 | 0.32 | 0.84 | 0.69 | 0.32 | 0.65 | 0.61 | 0.06 |

**Supplementary table D.** P-values for Spearman correlations between the 30 Bathyarchaeota OTUs and environmental parameters. Correlations were considered statistically significant when p-value<0.05. The physicochemical parameters are designated as SD (sediment depth), DO (dissolved oxygen), pH (potential of hydrogen), ORP (oxidation-reduction potential), EC (electric conductivity), N (total nitrogen), NH_4_^+^ (ammonium), NO_3_^-^ (nitrate), S (total sulphur), OM (total organic matter), OC (total organic carbon) and CH_4_ (methane).

| OTUs | SD | DO | pH | ORP | EC | N | NH<sub>4</sub><sup>+</sup> | NO<sub>3</sub><sup>-</sup> | S | OM | OC | CH<sub>4</sub> |
| --- | --- | --- | --- | --- | --- | --- | --- | --- | --- | --- | --- | --- |
| OTU7 | 0.03 | 0.21 | 0.78 | 0.26 | 0.35 | 0.53 | 0.91 | 0.29 | 0.74 | 0.23 | 0.27 | 0.01 |
| OTU12 | 0.03 | 0.18 | 0.96 | 0.18 | 0.18 | 0.57 | 0.65 | 0.42 | 0.82 | 0.29 | 0.35 | 0.03 |
| OTU22 | 0.32 | 0.91 | 0.21 | 0.78 | 0.65 | 0.74 | 0.75 | 0.16 | 0.65 | 0.01 | 0.02 | 0.09 |
| OTU26 | 0.03 | 0.49 | 0.82 | 0.57 | 0.61 | 0.42 | 0.51 | 0.16 | 0.53 | 0.26 | 0.33 | 0.02 |
| OTU41 | 0.1 | 0.42 | 0.35 | 0.35 | 0.39 | 0.82 | 0.73 | 0.29 | 0.78 | 0.03 | 0.04 | 0.01 |
| OTU64 | 0.53 | 0.8 | 0.59 | 0.84 | 0.45 | 0.19 | 0.66 | 0.15 | 0.49 | 0.13 | 0.14 | 0.4 |
| OTU76 | 0 | 0.04 | 0.69 | 0.13 | 0.32 | 0.43 | 0.82 | 0.73 | 0.53 | 0.65 | 0.7 | 0.02 |
| OTU86 | 0.23 | 0.69 | 0.71 | 0.82 | 0.57 | 0.16 | 0.97 | 0.11 | 0.38 | 0.19 | 0.23 | 0.19 |
| OTU100 | 0.07 | 0.02 | 0.6 | 0.11 | 0.25 | 0.31 | 0.68 | 0.82 | 0.37 | 0.77 | 0.77 | 0.17 |
| OTU107 | 0.42 | 0.93 | 0.63 | 0.93 | 0.53 | 0.22 | 0.9 | 0.06 | 0.57 | 0.15 | 0.18 | 0.3 |
| OTU128 | 0.33 | 0.11 | 0.06 | 0.29 | 0.53 | 0.55 | 0.71 | 0.45 | 0.87 | 0.12 | 0.14 | 0.87 |
| OTU160 | 0.03 | 0.35 | 0.91 | 0.46 | 0.49 | 0.35 | 0.73 | 0.18 | 0.57 | 0.32 | 0.38 | 0.03 |
| OTU167 | 0.41 | 0.27 | 0.1 | 0.4 | 0.82 | 0.98 | 0.91 | 0.29 | 0.91 | 0.14 | 0.17 | 0.87 |
| OTU208 | 0.66 | 0.69 | 0.43 | 0.87 | 0.61 | 0.61 | 0.14 | 0.63 | 1 | 0.06 | 0.1 | 0.33 |
| OTU212 | 0.22 | 0.86 | 0.69 | 0.73 | 0.41 | 0.44 | 0.56 | 0.09 | 0.73 | 0.11 | 0.16 | 0.13 |
| OTU215 | 0.23 | 0.63 | 1 | 0.84 | 0.53 | 0.08 | 0.93 | 0.16 | 0.32 | 0.38 | 0.46 | 0.29 |
| OTU361 | 0.38 | 0.09 | 0.25 | 0.02 | 0.15 | 0.13 | 0.62 | 0.08 | 0.25 | 0.56 | 0.62 | 0.52 |
| OTU373 | 0.53 | 0.27 | 0.01 | 0.57 | 0.61 | 0.3 | 0.98 | 0.38 | 0.61 | 0.03 | 0.04 | 0.69 |
| OTU381 | 0 | 0.14 | 0.26 | 0.34 | 0.63 | 0.34 | 0.47 | 0.79 | 0.39 | 0.72 | 0.66 | 0.08 |
| OTU388 | 0.03 | 0.09 | 0.91 | 0.26 | 0.69 | 0.53 | 0.55 | 0.53 | 0.78 | 0.91 | 0.89 | 0.06 |
| OTU456 | 0.03 | 0.12 | 0.93 | 0.24 | 0.47 | 0.49 | 0.81 | 0.4 | 0.76 | 0.53 | 0.56 | 0.03 |
| OTU622 | 0.41 | 0.93 | 0.42 | 0.71 | 0.35 | 0.84 | 0.56 | 0.1 | 0.91 | 0.03 | 0.05 | 0.16 |
| OTU783 | 0.32 | 0.35 | 0.87 | 0.23 | 0.06 | 0.57 | 0.95 | 0.49 | 0.96 | 0.12 | 0.15 | 0.23 |
| OTU3587 | 0.29 | 0.03 | 0.61 | 0.22 | 0.47 | 0.43 | 0.54 | 0.45 | 0.51 | 0.76 | 0.75 | 0.53 |
| OTU4026 | 0.45 | 0.05 | 0.53 | 0.03 | 0.07 | 0.74 | 0.53 | 0.49 | 0.91 | 0.12 | 0.11 | 0.23 |
| OTU4091 | 0 | 0.04 | 0.69 | 0.13 | 0.32 | 0.43 | 0.82 | 0.73 | 0.53 | 0.65 | 0.7 | 0.02 |
| OTU4395 | 0.01 | 0.08 | 0.86 | 0.09 | 0.22 | 0.77 | 0.82 | 0.56 | 1 | 0.48 | 0.52 | 0.01 |
| OTU5003 | 0.03 | 0.03 | 0.69 | 0.17 | 0.35 | 0.27 | 0.8 | 0.87 | 0.42 | 0.78 | 0.81 | 0.1 |
| OTU6022 | 0.64 | 0.59 | 0.11 | 0.8 | 1 | 0.93 | 0.68 | 0.11 | 0.78 | 0.01 | 0.02 | 0.22 |
| OTU9193 | 0.23 | 0.46 | 0.91 | 0.29 | 0.09 | 0.61 | 0.71 | 0.32 | 0.96 | 0.14 | 0.18 | 0.16 |

## 9 DECLARATIONS

## Authors’ contributions

- Conceived of or designed study: Pellizari V. H., Jovane L., Romano R. G.
- Performed research: Romano R. G., Bendia A. G., Franco D. C., Jovane L., Pellizari V. H.
- Analyzed data: Romano R. G., Franco D. C.,Bendia A. G., Signori C. N., Yu T., Wang F.
- Contributed new methods or models: Yu T., Wang F.

The first draft of the manuscript was written by Romano R. G. and all authors commented on previous versions of the manuscript. All authors read and approved the final manuscript.

## Data Availability

The datasets generated during the current study are available in the GenBank repository under BioProject ID PRJNA522377.

